# Milk-derived extracellular vesicles are functional mediators of postnatal neuroimmune rescue

**DOI:** 10.64898/2026.10.03.756449

**Authors:** Giovanna De Albuquerque Silva Braga, Penelope Pando, Hannah Hazzard, Hannah Vuong, Tran Tu Quyen Le, Yesim Ates, Serena B. Gumusoglu, Amanda C. Kentner

## Abstract

Early parental experiences can be translated into biological signals that are transmitted to offspring. These signals influence developmental trajectories to both increase offspring vulnerability following adverse experiences or promote resilience when environments are supported and enriched. Among the potential routes of parental-offspring communication, milk-derived extracellular vesicles (EVs) transferred through nursing are particularly interesting because they carry bioactive molecular cargos capable of influencing recipient cells. Previously, we found that milk-derived EV cargo is remodeled by both adverse and positive experiences and that environmental enrichment (EE) can rescue adversity-associated effects, including EV-miRNA cargo changes. Environmental programming of EVs during lactation may therefore have functional consequences on offspring development and adaptability. Here, we tested whether milk-derived EVs from our rat EE model are sufficient to rescue offspring outcomes following maternal immune activation (MIA). Milk-derived EVs isolated from donor EE dams were administered directly to MIA-exposed pups, mitigating MIA-associated reductions in social behavior. EE-EV administration also promoted social behavior in non-MIA exposed male offspring. To investigate microglia as a cellular target, EE-EVs were administered to LPS-activated BV2 microglial cells. EE-EV exposure attenuated LPS-associated transcriptional alterations across inflammatory pathways, including cytokine, chemokine, and NF-ĸB signaling, while modulating genes associated with cellular homeostasis and metabolic responses. Together, these results 1) support milk-derived EVs as a biologically active interface through which environmental experience may be communicated to offspring, and 2) identify microglial transcriptional responses as a potential cellular pathway underlying these effects.

## Introduction

Parental stress, inflammation, and nutrition, alongside caregiving behaviors represent key environmental signals that can shape offspring developmental trajectories and influence the maturation and function of endocrine, immune, and behavioral systems (Bell and Hellmann, 2019; Schneider et al., 2024; Ribas et al., 2024; Kentner et al., 2019; Chan et al., 2018; Pipelias et al., 2026; Wijenayake et al., 2023). The consequences of these signals are not uniformly detrimental; rather, developmental trajectories may be shaped by the nature and context of parental experiences (McDorman et al., 2024; Kentner et al., 2019; Schneider et al., 2024; Neylan et al., 2025). For example, adverse experiences can increase susceptibility to maladaptive outcomes (Ribas et al., 2024; Pipelias et al., 2026), while supportive and enriched environments can promote neuroplasticity and adaptive responses, fostering resilience even in times of adversity (Kentner et al., 2019). However, the ways in which these diverse experiences are translated into biological signals and conveyed to the developing offspring remain poorly understood. In particular, it remains unclear whether distinct experiences engage shared or experience-specific signals and, importantly, which molecular carriers transmit this information.

Among the potential molecular mediators of parental-offspring communication, the transfer of milk-derived extracellular vesicles (EVs) through breastfeeding/nursing is of particular interest. EVs are membrane-bound particles that serve as endocrine signaling factors, carrying bioactive cargos including microRNAs (miRNAs), proteins, lipids, and other regulatory molecules which would otherwise be degraded (Welsh et al., 2024; Jiang et al., 2021; Kalluri et al., 2020). Milk-derived EVs are well positioned for this role because their cargo can withstand gastrointestinal conditions and are taken up by intestinal cells, thereby providing a route for these putative lactational signals to enter the body (Liao et al., 2017; Khan et al., 2018).

Animal studies further demonstrate that milk-derived EVs can distribute beyond the gastrointestinal tract, reaching peripheral tissues and the brain following experimental administration or natural transfer during nursing (Manca et al., 2018; López de Las Hazas et al., 2022; Zhou et al., 2022; Haney et al., 2019). This bioavailability is especially relevant during early development in rodents when some brain associated barriers are more permeable to small molecules (Saunders et al., 1999; Haney et al., 2019).

Milk-derived EVs also interact with and modulate neural (Gönüllü et al., 2026; Storm et al., 2025; Zhou et al., 2022) and immune cell populations, including microglia (Akinduro et al., 2025; Storm et al., 2025; Wijenayake et al., 2024; Li et al., 2026). Microglia provide a dynamic cellular connection between peripheral signals and neuroimmune function, responding to systemic and local cues through transcriptional programs that regulate immune responses, cytokine signaling, and cellular homeostasis (Di Pietro et al., 2026). Moreover, they respond to inflammatory stimuli, including LPS-programmed EVs (La Torre et al., 2022; Santiago et al., 2023). Indeed, EVs themselves are responsive to inflammatory state; LPS exposure alters the quantity and protein composition of EVs released by BV2 microglia, including their cytokine cargos (La Torre et al., 2022), and the miRNA profile of milk-derived EVs (Martz et al., 2026). These findings underscore the view that milk-derived EVs and their unique molecular cargo signatures reflect information about the environment from which they are produced, transmitting this information to recipient cells (La Torre et al., 2022; Wijenayake et al., 2023).

In this context, the lactational period may provide more than nutritional support; it may represent a functional route through which information about the parental environment is conveyed to the developing offspring (Wijenayake et al., 2023). Supporting this possibility, our recent work demonstrated that environmental conditions during lactation alter the miRNA cargo of milk-derived EVs, matching miRNA changes in the hippocampus of nursing rodent pups (Martz et al., 2026). Importantly, housing nursing litters in environmental enrichment (EE) buffered or restored maternal immune activation (MIA)-associated changes in both the milk-derived EV and offspring hippocampal miRNA profiles, in addition to rescuing the social behavior of offspring. These findings established that milk-derived EV cargos are dynamically responsive to both adverse and enriched environmental conditions.

Emerging evidence suggests that EVs can function as biologically active mediators rather than simply reflect physiological states. For example, a recent study demonstrated that circulating EVs improves sociability in mice with genetically associated social deficits via EV-miRNAs, with effects on neuronal signaling in the prefrontal cortex (Matoba et al., 2026). The biological activity of EVs is further supported by studies showing that they can convey environmentally derived information across reproductive and developmental contexts. Reproductive tract EVs, for example, transmit paternal environmental information to influence offspring neurodevelopment and stress-related phenotypes (Chan et al., 2020) while placental EVs have been implicated in maternal-fetal communication to influence development (see Hazzard et al., 2026). Together, these findings support a model in which milk-derived EVs could mediate communication between parental peripheral experiences and offspring developing tissues, including the brain.

In the present study, we extend our previous findings to test whether milk-derived EVs directly convey environmental enrichment signals to offspring. We isolated milk-derived EVs from EE-housed donor dams and administered them directly to unrelated offspring reared by either MIA-exposed or unexposed control dams. This design separates the potential contributions of EE-programmed milk from those of the maternal environment (e.g., maternal care, housing). We hypothesized that EE-programmed EVs would attenuate MIA-associated deficits in social behavior while promoting social behavior in unexposed control offspring. Moreover, given the role of microglia in responding to MIA and in sculpting brain circuits involved in sociability, we examined cellular mechanisms that could link EV exposure to microglial function. Using the immortalized BV2 microglia cell culture model (Yang et al., 2024), we examined whether EE-EVs modify transcriptional responses to an inflammatory challenge. Together, these experiments test a mechanistic model in which environmental enrichment alters milk-derived EV signals, promoting offspring resilience and modulating neuroinflammatory and homeostatic responses to MIA.

## Methods and Experimental Overview

### Animals

Female Sprague Dawley rats (Charles River, Wilmington, MA) were housed at 20°C on a 12/12 h light/dark cycle (6 A.M. to 6 P.M.) with *ad libitum* access to food and water. Animals were reared in either environmental enrichment (EE; 91.5 × 64 × 159 cm sized multi-level cages with multiple ramps, tubes, chew bones, and toys) or standard sized laboratory cages (SD; 27 × 48 × 20 cm, with a single tube and a chew bone), as described in our previous work (Connors et al., 2014; Connors et al., 2015; Núñez Estevez et al., 2020; DeRosa et al., 2022a; Martz et al., 2026). Both caging conditions had access to Bed-r’Nest® nesting materials and animals were housed in pairs. Male breeders were pair-housed in SD cages unless they were breeding, which took place one week after animals were placed into their assigned housing conditions. The breeding configuration was two females and one male in either an EE cage, or a larger SD one-level cage (51 × 41 × 22 cm, with a single tube and a chew bone). At approximately gestational day 16, dams were separated and individually housed in their assigned condition, with tubes and toys removed to prevent pup injury, as described previously (Connors et al., 2014; Connors et al., 2015; Núñez Estevez et al., 2020; DeRosa et al., 2022a; Martz et al., 2026). Day of birth was designated as postnatal day (P)0. All animal protocols were approved by the Massachusetts College of Pharmacy and Health Sciences (MCPHS) Institutional Care and Use Committee and were carried out in compliance with the Association for Assessment and Accreditation of Laboratory Animal Care.

### Milk collection from donor dams

EE-housed donor dams were separated from their litter for 2 hours, to allow milk to accumulate for sample collection, as previously described (DeRosa et al., 2022a; DeRosa et al., 2022b; Martz et al., 2025). Donor dams were anesthetized with isoflurane in O2 and administered 0.2 mL of oxytocin (20 USP/mL i.p.) to facilitate milk production (Paul et al., 2015). Given the pharmacokinetic profiles of these drugs, nursing can resume immediately after the milking procedure as neither isoflurane nor oxytocin are absorbed by the feeding offspring (Drugs and Lactation Database, 2020; Lee & Rubin, 1993; Par Pharmaceutical Inc, 2020). After cleaning the teats with distilled water, milk was collected by gently squeezing the base of each teat and collecting the milk, until the required sample volume was obtained (∼45-60 minutes of collection time to acquire ∼2.0 mL sample volume). Milk was collected from dams between P1 and P11, with no more than three collection points per dam, and later pooled during the EV isolation process (see details below). This timing accounts for two critical nursing stages: early lactation, when colostrum has relatively higher concentration of EVs and mid-lactation when milk is mature and yields higher sample volumes (Ventrella et al., 2021; Samuel et al., 2017). Once dams fully awoke from the procedure they were returned to their litters in their home cage. During the separation and milk collection period, pups from each litter were housed together on heating pads in a smaller cage to keep them warm (see DeRosa et al., 2022a; DeRosa et al., 2022b; Martz et al., 2026). Dams and their litters were euthanized by isoflurane overdose at the end of the milk collection protocol at P11. No offspring from any milk donor litters were evaluated on any study measures.

### Isolation and characterization of milk-derived extracellular vesicles (EVs) from donor dams

Isolation and characterization of milk-derived extracellular vesicles (EVs) from EE donor dams is outlined in **Figure 1** and followed the procedures previously described (Martz et al., 2026). Briefly, milk collected from each donor dam was centrifuged at 3,000 x g for 10 min at room temperature (RT), to separate out the cream layer. The supernatant was removed and placed into a fresh tube and centrifuged again at 3,000 x g for 10 min RT. The cream layer was removed and the clear supernatant stored at −80°C until further processing.

**Figure 1.**
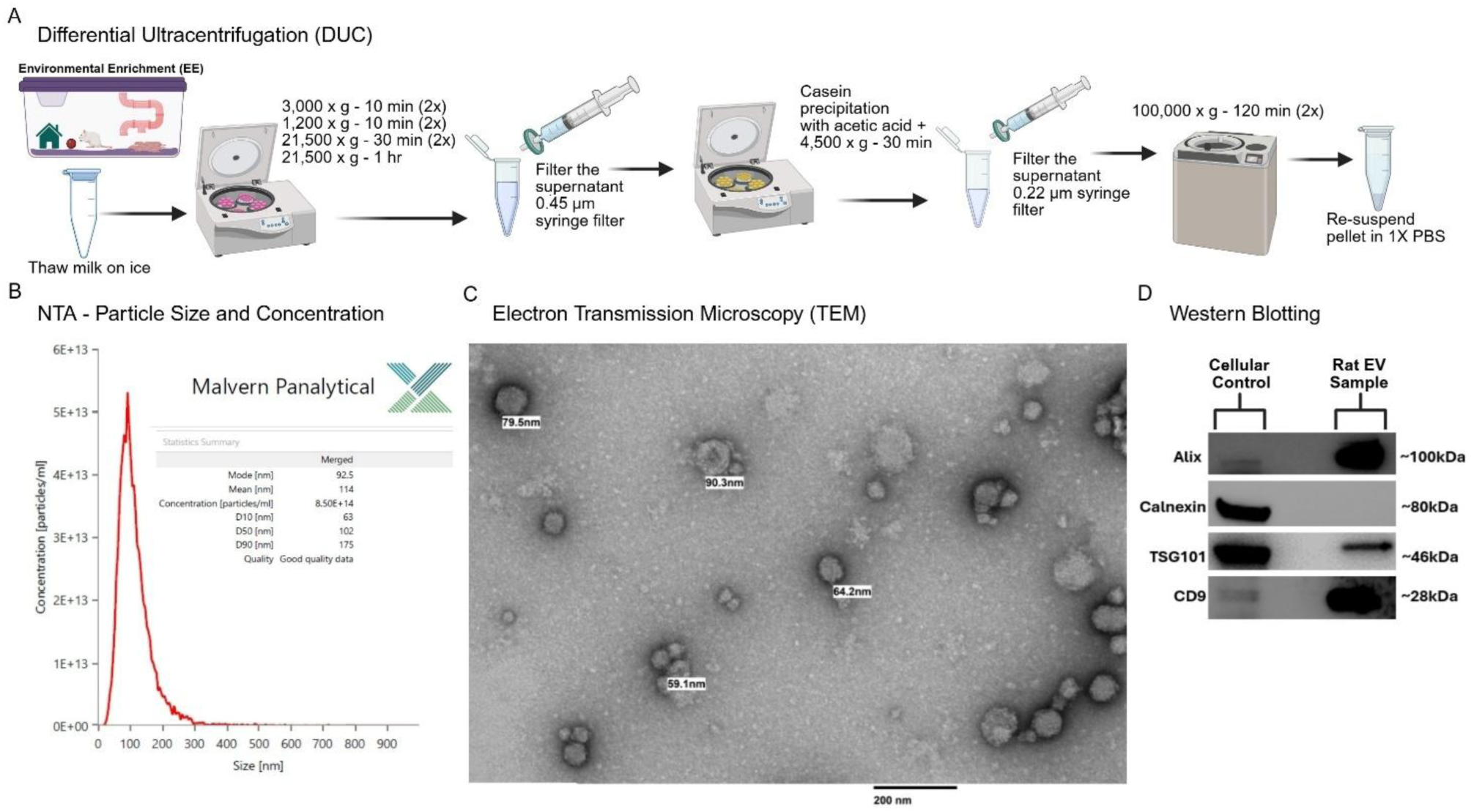
Characterization of milk-derived extracellular vesicles (EVs) isolated from environmentally enriched (EE) donor dams. (**A**) The differential ultracentrifugation (DUC) and serial filtration protocol followed for EV isolation. (**B**) Size (nm) and concentration (particles/mL) of EVs from EE donor dams, as determined by nanoparticle tracking analysis (NTA) with a mean size of 114 nm. (**C**) Transmission electron microscopy (TEM) of EVs isolated from EE donor dams. Scale bar is 200 nm with some individual EE-EVs identified in sizes ranging from 59.1 to 90.3 nm. (**D**) The presence or absence of positive (Alix, TSG101, CD9) and negative (Calnexin) protein markers for EE-EVs as determined by western immunoblotting. *Created in BioRender. Kentner, A. (2026)* https://BioRender.com/w09n833.

Differential ultracentrifugation, combined with serial filtration (DUC), is identified in the Minimal Information for Studies of Extracellular Vesicles (MISEV) 2023 guidelines as an acceptable method for extracellular vesicle isolation (Welsh et al., 2024). Indeed, DUC yields high-quality MEV concentrations appropriate for biological experiments (Wijenayake et al., 2021; Wijenayake et al., 2024; Martz et al., 2026). For this approach (**Figure 1A**; Martz et al., 2026), samples were thawed on ice and centrifuged twice at 3,000 x g for 10 min at 4°C. The supernatants were then centrifuged at 1,200 x g for 10 min at 4°C (2x), at 21,500 x g for 30 min at 4°C (2x), and once at 21,500 x g for 1 h at 4°C. To clear fat globules and cellular debris, samples underwent filtration through 0.45 µm PES syringe filters (Ultident; 229749). Casein precipitation, via 17.5 molar acetic acid (1:100 (v/v)) added to each sample Morozumi et al., 2021), was followed by centrifugation at 4,500 x g for 30 min at 4°C. To remove carry over precipitates and remaining cell debris, the supernatant was filtered through 0.22 µm PES syringe filters (Ultident; 229747). Following this, the supernatants (now called the whey fraction) were centrifuged at 100,000 x g for 120 minutes (2x) using a Beckman Coulter XPN100 ultracentrifuge and SW55 TI (Beckman Coulter) swing bucket at 4°C. The pellets containing EVs were resuspended in filtered 1X phosphate buffer solution (PBS) using a 1:2 (v/v of filtered PBS to whey fraction; Martz et al., 2026).

During isolation, EV samples were allocated for downstream analyses and experimental applications. A subset of individual EV aliquots was reserved for transmission electron microscopy (TEM), nanoparticle tracking analysis (NTA), and western blotting, to further validate EV preparations, according to MISEV 2023 guidelines (described below; Welsh et al., 2024) as we have done previously (Martz et al., 2026).

The biological unit of study was considered the offspring and as such EV samples (n = 8 donor dams) were pooled to generate a single standardized master EE-EV stock. Stock was then aliquoted for individual treatments and stored at −80°C until neonatal administration (see section on neonatal offspring EE-EV treatments) or BV2 microglia experiments.

#### NTA

The concentration and particle size of EVs was immediately determined using light scattering based nanoparticle tracking analysis with a NanoSight Pro (Malvern Instruments, Ltd.) following the manufacturer’s instructions. Samples were diluted with filtered 1x PBS. Data capture settings were adjusted to a flow rate of 4.0 µl/min with five replicates of 17 second captures using a Green 532 nm laser; data show a high particle concentration and a mean EV particle size of 114 nm (**Figure 1B**).

#### TEMS

EV suspensions were diluted 1:10 (v/v) in 1X PBS. 5 uL samples were each absorbed for 1 minute to a carbon coated grid (CF400-CU, Electron Microscopy Sciences) that had been made hydrophilic by glow discharge at 25 mA for 20 seconds. Whatman #1 filter paper was used to remove excess liquid, and the grid was briefly floated on a drop of double distilled water (ddH2O) to remove residual salts or phosphate, blotted three times on filter paper, and negatively stained with 1% uranyl acetate (22400, Electron Microscopy Sciences) for 20-30 seconds at RT. To verify EV morphology, images were acquired at the Harvard Electron Microscopy Core Facility using a Tecnai™ G² Spirit BioTWIN transmission electron microscope equipped with an AMT Nanosprint 43-MKII camera (**Figure 1C**).

#### Western blotting

Following extraction, EV protein concentrations were measured using a Pierce™ BCA assay (Thermo Scientific, 23227) to determine protein loading amounts. EE-EVs were validated by the presence of the following positive EV markers: Alix (HY-P80011, MCE, 1:1000), TSG101 (Ab125011, Abcam, 1:1000), and CD9 (E607008, Booster, 1:1000), in addition to the absence of Calnexin (12186, Signalway Antibody [SAB], 1:1000), which was used as a negative protein marker. Protein samples and Hep G2 cells (cellular control) were loaded (50 µg) and run on 12% SDS-polyacrylamide gels which were then transferred to a PVDF membrane (Millipore, IPVH00010). Membranes were then blocked with 5% skim milk in TBST for 1 hour. After washing in 1X TBST (3 x 10 minutes), primary antibodies were incubated at 4°C overnight. Membranes were again washed 3 x 10 minutes in TBST and incubated in secondary antibody (31460, Invitrogen, 1:5:000) at room temperature for 1 hr. After washing (as described above), membranes were visualized using ECL Plus Chemiluminescence Reagent (RPN2232, Cytiva). Western immunoblotting confirmed the presence of markers CD9, Alix, TSG101, and the absence of Calnexin (**Figure 1D**). Protein markers were not quantified but rather used to determine whether they were absent or present in EE-EVs and cellular controls to validate EV samples (Wijenayake et al., 2021; Wijenayake et al., 2024; Welsh et al., 2024; Martz et al., 2026).

### Maternal immune activation (MIA) induction and neonatal offspring EE-EV treatments

The timeline of the general animal procedures is outlined in **Figure 2A**. Briefly, Sprague Dawley female rats (Charles River, Wilmington) were bred and reared in SD housing. Across three separate cohorts, dams were treated intraperitoneally (i.p) with either 100 µg/kg lipopolysaccharide (LPS; Escherichia coli, serotype 026:B6; L-3755; n = 12) or equivolume pyrogen-free saline (Saline; n = 11) on P10 (DeRosa et al., 2022; Martz et al., 2025; Kentner et al., 2019: **Supplemental Table 1)**. Simultaneously, to test whether EE-EVs were protective, male and female pups from the LPS litters were randomly assigned to receive either PBS (LPS+PBS; control), EVs derived from EE donor dams (LPS+EE-EV), or handling (LPS+Handled) using a within litter treatment design to control for maternal care. To counter the lower bioavailability associated with oral EV administration (Haney et al., 2019; Manca et al., 2018; Wiklander et al., 2015), on P10, LPS+EE-EV pups received (9 x 10^11^ EV particles/gram of body weight per injection) via bilateral i.p injections (left and right side; total dose of 1.80 x 10^12^ EV particles/gram of body weight per treatment day) or an equivalent volume of PBS using the same injection protocol (LPS+PBS group).

**Figure 2.**
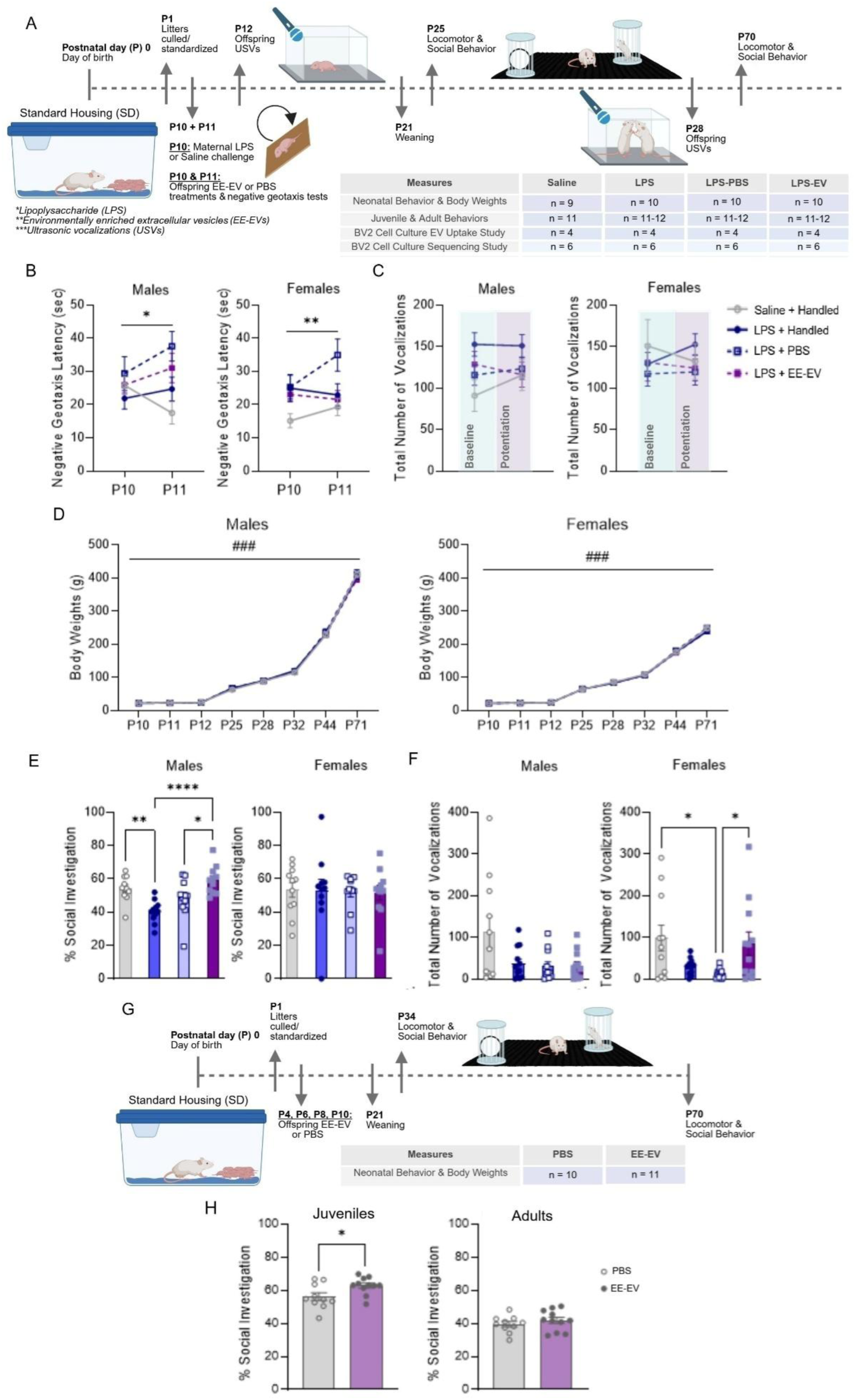
Milk-derived extracellular vesicles (EVs) from environmentally enriched (EE) donor dams both rescue and promote social behavior. **(A)** Timeline of experimental procedures and group designations/sample sizes (n) to test whether neonatal administration of EE-EVs protects against maternal immune activation (MIA)-induced alterations to offspring behavior. (**B**) Negative geotaxis performance across postnatal days (P) 10 and 11 was delayed in LPS+PBS male (left) and female (right) offspring. Neither (**C**) total number of P12 isolation-induced vocalizations emitted nor (**D**) body weight gain (g) was affected in male (left) or female (right) offspring following MIA. Data expressed as mean ± SEM, n = 9-10 litters/group. *p <0.05, **p <0.01; Saline+Handled versus LPS+PBS; ###p<0.001, main effect of time. Juvenile behavioral outcomes are shown for males (left) and females (right) for the (**E**) percent time investigating a novel conspecific on P25 and the (**F**) total number of play vocalizations emitted on P28. (**G**) Timeline of experimental procedures and group designations/sample sizes (n) to test whether neonatal administration of EE-EVs promotes social behavior in healthy rat offspring. Juvenile (left) and adult (right) behavioral outcomes are shown for male offspring (**H**) percent time investigating a novel conspecific on P34. Data expressed as mean ± SEM, n = 10-12 litters/group. Comparisons shown indicate *p <0.05, **p <0.01, ***p <0.001, ****p <0.0001. LPS: lipopolysaccharide; EE: environmental enrichment; EV: extracellular vesicles. *Created in BioRender. Kentner, A. (2026)* https://BioRender.com/t61q425.

Handled pups from the Saline (Saline+Handled) and LPS litters were gently picked up and received a light finger poke on each of their right and left abdomens with the experimenter’s pointer finger, to mimic the injection procedure of the LPS+PBS and LPS+EE-EV groups. After administration, pups and dams were immediately reunited in their home cages. These procedures (adapted from Drommelschmidt et al., 2017) were repeated 24-hours later on P11.

In a separate experiment, to test whether EE-EVs promote social behavior independently in healthy animals, male pups from SD reared mothers (untreated) received bilateral i.p injections (2 x 10^10^ EV particles/gram of body weight per injection; left and right side; total dose of 4 x 10^10^ EV particles/gram of body weight per treatment day) or an equivalent volume of PBS using the same injection protocol as above on P4, P6, P8, and P10.

All offspring were weaned on P21 and continued to be SD-housed. Each experimental group included only one pup per litter and sex. No litters from the EE-EV administration experiments underwent milking or related separation procedures.

### Offspring Behavioral Measures

To test whether EE-EVs have functional effects *in vivo*, male and female offspring were evaluated on a battery of behavioral tests across P10 through to adulthood. Behavioral assessments were counterbalanced and evaluated by observers’ blind to group designations.

#### Negative geotaxis (P10 & P11)

Four hours following each of the P10 and P11 treatments, offspring were positioned facing downward on a 45° slanted surface and their latency to complete a 180° turn was recorded. This sensorimotor task evaluates the natural developing tendency of rodents to turn upright on a slope, in response to vestibular gravity cues (Ruhela et al., 2019; Feather-Schussler and Ferguson, 2016). Each trial ended if the pup had not yet fully rotated within 60 seconds. Animals completed three successive trials, and the average rotation time was used for analysis. Using two apparatuses, this test took ∼15 minutes per litter to complete. During this period, each dam was kept in a separate “vacation” cage while her litter was being evaluated. Her pups were held together in a smaller cage on a heating pad until each experimental pup was evaluated.

#### Ultrasonic vocalizations (USVs; P12 & P28)

On P12, pups were individually transferred to a small cage lined with Nestlets®. To assess maternal potentiation of pup USVs, this cage was placed inside a sound-attenuating chamber, and isolation calls (negative valence) were recorded using an ultrasonic microphone (UltraSoundGate 116-200) positioned 20 cm above the floor (baseline phase). Following a brief reunion (5 minutes) with their dams in the home cage, each pup was separated again and retested under the same conditions (potentiation phase; adapted from Bodi et al., 2016; Martz et al., 2023; Merengueli et al., 2025). On P28, offspring were placed with an unfamiliar conspecific from the same treatment condition and social play vocalizations (positive valence) were recorded for five minutes as described above (Martz et al., 2023). The USV microphone was connected to a computer and the vocalizations captured using Audacity at a sampling rate of 192 kHz, to capture a range of USVs. Total number of vocalizations was analyzed by the automated LMT USV analysis software (de Chaumont et al., 2019; Merengueli et al., 2025).

#### Locomotor activity and social behavior (P25 and P70)

Open field and social preference tests were video recorded and evaluated in juvenile and adult animals as previously described (see Yan & Kentner, 2017; Connors et al., 2014; Zhao et al., 2021a; Núñez Estevez et al., 2020; Martz et al., 2023; Martz et al., 2026). For open field, distance travelled (m) and time (seconds) spent in each of the perimeter and center of the field was analyzed using automated behavioral monitoring software (Any-maze, Wood Dale, IL) and data presented as a percentage of time spent in the center of the open field [(total time in center/total time in perimeter + total time in center) × 100] as an index of anxiety-like behavior.

Following the five-minute open field habituation period, two cleaned wire containment cups were placed on each end of the arena for a five-minute social preference test. One cup held a novel untreated animal of the same sex, age, size, and strain and the second cup held a novel object. The locations of the novel rats (n =6) and objects were interchanged between trials. An animal was actively investigating when its head was within 7 cm of a containment cup, or it was touching the cup. Data are presented as a percentage of time spent in social investigation [(total time with novel animals/total time with novel object) + (total time with novel animal) × 100] across a five-minute period.

### BV2 microglial cell culture, EE-EV treatment, and RNA sequencing

To investigate potential molecular mechanisms underlying behavioral rescue by EE-EVs, the immortalized BV2 female murine microglial cell line (gift of the Dailey Laboratory, The University of Iowa; Eyo et al., 2013) was grown in 6-well plates containing Dulbecco’s modified Eagles’ medium (DMEM) and supplemented with 10% fetal bovine serum (FBS, Gibco, Grand Island, NY, USA) and 1% penicillin-streptomycin (Gilbo). The cells were maintained in an incubator (Sanyo) at 37°C with 5% CO2. Cells were counted with a hemocytometer (Curiosis, cat# CRCSD-NI50) and plated into 24-well plastic culture plates to culture at a density of 3 x 10^5^ cells per well (adapted from Yang et al., 2024). Cells used in the experiment were between passage numbers 22 and 32. Cell morphology and overall health were routinely monitored to confirm expected BV2 phenotype and no visible signs of cellular stress, abnormal growth, or death.

To confirm EV interaction with recipient microglia, EVs were brought to room temperature and dyed with the lipophilic fluorescent dye PKH-26 (PKH; Sigma-Aldrich, St Louis, MO, USA). EVs were incubated with 1% PKH in dilutant C for 5 minutes before being quenched with 1.2x the volume of 10% BSA. To further deplete unbound dye, EVs were re-isolated using the Izon Automated Fraction Collection with qEV 35nm columns on default settings. EV-enriched fractions 1 and 2 were pooled and used for EV treatment as described by the manufacturer (<u>IZON qEVORIGINAL User Manual - GEN 2</u>, https://www.izon.com/resources). BV2 cells were then cocultured with either 10^8^, 10^9^, or 10^10^ EV particles/mL (n = 4 wells per concentration) for 12 hours to determine the optimal EV concentration for uptake. Following incubation, BV2 cells were detached from the plate using 0.25% Trypsin-EDTA (Gilbco, cat# 25200056) before being pelleted (1000 x g, 3 minutes) and washed with PBS/1% BSA. Cells were filtered at 40µm and stained with Fc block (1:50; Invitrogen, cat# 56-0451-82) for 5 minutes followed by CD45/AF700 stain (1:50; Invitrogen, cat# 56-0451-82) for 20 minutes. EV uptake was examined with the Amnis ImageStream using the following laser settings: 405nm: 120mW, 488nm: 100mW, 561nm: 150mW, 642nm: 150mW, 785nm: 3.59mW at 40 x magnification.

To evaluate inflammatory responses, a different set of cells were plated and serum starved overnight to align cell cycles prior to treatment with either LPS (100 ng/mL; *E. coli* K-12, InvivoGen, cat# tlrl-eklps; Roosen et al., 2021) or PBS every 12 hours (for a total of four treatments). On the last treatment, EVs (10^10^ EV particles/mL) or PBS were added to each well across two replicates and then 12 hours later cells were collected and RNA extracted using the Qiagen RNeasy Plus Mini Kit, according to the manufacturer’s instructions (Qiagen, 74134). The cell culture groups were 1) PBS+PBS, 2) LPS+PBS, 3) PBS+EE-EV, and 4) LPS+EE-EV (n = 6 wells/group). EV concentrations were confirmed prior to cell culture experiments using the Horiba Viewsizer 3000 (Horiba, Austin, TX) with the following parameters: laser power blue 210mW, green 12mW, red 8mW, gain 30dB, exposure 15ms, recording 25 videos at 150 frames/video, integration range 30-500nm. Samples were diluted 1:1,000-1:10,000 in 0.22µm filtered 1 x PBS which was also used for blank-subtraction.

RNA sequencing (RNA seq) was performed by Plasmidsaurus, Inc (Kentucky, USA) using a 3′-end counting RNA-seq workflow and an Illumina NovaSeq X Plus instrument. RNA was converted to complementary DNA (cDNA) using a poly(dT)VN primer, followed by second-strand synthesis, tagmentation, library indexing, and amplification. Libraries were sequenced using stranded, single-end sequencing on an Illumina NovaSeq X Plus platform.

Raw FASTQ files were assessed for quality using FastQC v0.12.1 and quality filtered using fastp v0.24.0, including poly-X tail trimming, 3′ quality-based tail trimming, a minimum Phred quality score of 15, and a minimum read length of 50 bp. Quality-filtered reads were aligned to the *Mus musculus* (GRCm38) reference genome using STAR v2.7.11 with non-canonical splice junction removal and output of unmapped reads. Aligned reads were coordinate sorted using SAMtools v1.22.1. PCR and optical duplicates were removed using unique molecular identifier (UMI)-based deduplication with UMIcollapse v1.1.0. Alignment quality, strand specificity, and read distribution across genomic features were assessed using RSeQC v5.0.4 and Qualimap v2.3, with quality-control results summarized using MultiQC v1.32. Gene-level expression was quantified using FeatureCounts (Subread v2.1.1) with strand-specific counting, fractional assignment of multimapping reads, and exons and 3′ untranslated regions (UTRs) as feature identifiers, with reads grouped by gene_id. Gene-level counts were annotated using gene biotype and additional metadata obtained from the reference GTF file. Counts were normalized using the trimmed mean of M-values (TMM) method.

### Statistical analyses

Graph Pad Prism (version 11.0.1) was used to perform statistical analyses on behavioral data and for the BV2 microglial EV uptake study. While both male and female animals were included in the MIA rescue experiments, separate analyses were run for each sex (Martz et al., 2026; Merengueli et al., 2025). Previous cross-fostering work has associated nursing by EE-housed dams to increase social behavior in healthy male rats, but not females (DeRosa et al., 2022), therefore, we studied males in the EE-EV sociability experiment. Repeated two-way ANOVA (group by time), one-way ANOVAs, or t-tests were run as appropriate. Because the data was not normally distributed for the male and female P28 social USVs, the male P25 percent time spent investigating the novel object, or the male P70 percent time spent in social investigation measures, these statistics were analyzed on log transformed data. For clarity, the data figures display non-transformed data as mean ± SEM, and Tukey post hoc tests were used to account for multiple comparisons.

For the BV2 cell culture sequencing experiment, differential expression analysis was performed via the Plasmidsaurus RNA-seq analysis pipeline using edgePython (edgePy) to evaluate gene-level expression differences between user-defined sample groups in pairwise comparisons. Statistical significance was assessed using the false discovery rate (FDR) reported by Plasmidsaurus. Genes were considered differentially expressed when they demonstrated an absolute log2 fold change (|log2FC|) ≥2 and an FDR ≤0.01. Those with a log2FC ≥2 were classified as upregulated, whereas genes with a log2FC ≤−2 were classified as downregulated. Differentially expressed gene (DEG) lists were subsequently used for downstream functional and pathway enrichment analyses. Volcano plots for each comparison were constructed using the log2FC and-log10 (FDR) of expression in GraphPad Prism. Heatmaps representing our expression data were created using Morpheus (https://software.broadinstitute.org/morpheus).

To visualize transcriptional rescue, log2FC from the PBS+PBS versus LPS+PBS comparison (x-axis) were plotted against log2FC from the PBS+PBS versus LPS+EE-EV comparison (y-axis) for LPS-responsive genes. Genes showing movement toward a log2FC of zero following EE-EV treatment were considered consistent with transcriptional reversal (adapted from Kelly et al., 2019; Theisen et al., 2019). Counts per million (CPM)-based ANOVA and Tukey’s post hoc were applied to genes differentially expressed under LPS+PBS that returned toward PBS+PBS levels by LPS+EE-EV co-treatment. Additionally, these genes were split by the direction of the LPS effect (up versus down regulated) and tested for GO biological process overrepresentation in PANTHER 19.0 (Fisher’s exact test, Benjamini-Hochberg FDR; GO release 2026-04-28). The reference list was the set of genes detected in this RNA-seq dataset (13,055 Ensembl gene IDs, of which 9,733 mapped to GO-annotated mouse genes). Enriched terms (FDR < 0.05) were reduced to a non-redundant set by retaining only terms with no enriched GO descendant and plotted in R 4.3 with ggplot2.

Candidate miRNA–mRNA relationships were evaluated by integrating miRNAs we previously identified in milk-derived EE-EVs (Martz et al., 2026; GEO Accession Number GSE300818) with DEGs identified in the BV2 cells of the present RNA-seq study. Predicted miRNA targets were obtained from miRDB.org. While some lower target score candidates were evaluated, interactions with a target score >80 were considered high-confidence candidates. Predicted targets were then compared with LPS-responsive DEGs and their CMP-based ANOVA expression following EE-EV treatment to identify candidate miRNA–mRNA relationships associated with transcriptional rescue.

## Results

### EE-derived EVs are associated with attenuation of an early sensorimotor phenotype following MIA

To evaluate the immediate effects of MIA and EE-EV treatment, neonatal offspring behavior was assessed. Two-way repeated measures ANOVAs (males: F(3, 35) = 3.311, p = 0.0312; females: F(3, 35) = 3.943, p = 0.0159) showed significant group effects on negative geotaxis performance. Both male (p = 0.0389) and female (p = 0.0089) LPS+PBS exposed neonates required more time than Saline+Handled controls to reorient themselves upward on the inclined plane (**Figure 2B**). Although the LPS+PBS and LPS+EE-EV animals did not differ from one another (p > 0.05) the absence of group differences between LPS+EE-EV versus Saline+Handled, and LPS+Handled animals indicate a partial EE-EV-associated attenuation of the MIA-associated sensorimotor phenotype that emerged following the additional neonatal injection experience. Moreover, the timing of this attenuation corresponds to when milk EVs reportedly reach the brain (Khanam et al., 2023; Manca et al., 2018). In contrast, there were no significant group differences with respect to P12 isolation induced USVs (**Figure 2C**) or body weight gain (**Figure 2D**) across the study (p >0.05).

### EVs from EE donor dams rescue social behavior following MIA

For male juvenile offspring, significant group effects emerged for the percent of time spent in social investigation (F(3,40) = 9.099, p = 0.0001): LPS+Handled males engaged in less social investigation than Saline+Handled animals (p = 0.0042). Notably, neonatal treatment with EVs derived from EE donor dams reversed this effect (LPS+Handled versus LPS+EE-EV: p = 0.0001; LPS+PBS versus LPS+EE-EV: p = 0.032; **Figure 2E**), increasing social investigation levels (female offspring: p>0.05). In contrast, group effects were observed in female (F(3, 39) = 3.649, p = 0.0207), but not male (p>0.05), offspring for the number of P28 social play vocalizations emitted, with Tukey post hoc tests indicating that LPS+PBS females produced fewer vocalizations than both Saline+Handled (p = 0.0273) and LPS+EE-EV (p = 0.0415) groups (**Figure 2F**).

Adult LPS+EE-EV male animals engaged in significantly more social investigation than LPS+Handled males (p = 0.0224; **Supplemental Figure 1A**), highlighting the affiliative promoting effects of the EE donor EVs.

Across juvenile and adult groups, there were no significant differences in the percentage of time spent in the center of an open field or in total distance traveled (p>0.05; **Supplemental Figure 1B-E**), indicating that the observed social behavior changes were not driven by anxiety-like or locomotor effects.

### EVs from EE donor dams promoted social behavior in healthy offspring

To evaluate whether EE-EVs influence social behavior independently of MIA, we administered them to healthy offspring from untreated SD dams. This approach allowed us to isolate a transferable component of the lactational EE experience and test whether milk-derived EVs could carry signals associated with enrichment to offspring, without the potential confound of “enrichment loss” that occurs when animals are removed from EE (Smith et al., 2017). While neonatal EE-EV administration had no effect on negative geotaxis, USVs, nor open field activity (p >0.05, **Supplemental Figure 2A-E**), EE-EV treatment modestly augmented social investigation in juvenile, but not adult, male offspring (juveniles: t(19) = 2.378, p = 0.028; adults: p>0.05, **Figure 2G-H**).

### EE-EVs promote transcriptional rescue from LPS-induced inflammatory activation in BV2 microglia

To determine whether EE-EVs modulate the inflammatory response of microglia, we examined the transcriptional effects of this treatment on LPS-stimulated BV2 cells. First, EV uptake was established to identify the optimal concentration for the sequencing study. BV2 cells were exposed to three concentrations of PKH-dyed EVs, and uptake was quantified by cytometry (**Figure 3A**; **Supplemental Figure 3**). EV uptake increased in a concentration-dependent manner (F(2, 9) = 5884, p = 0.0001), with 10^10^ particles/mL demonstrating the greatest uptake compared to 10^8^ (p = 0.0001) and 10^9^ particles/mL (p = 0.0001). Therefore, 10^10^ EV particles/mL were selected for subsequent experiments to maximize EV exposure and uptake by BV2 cells.

**Figure 3.**
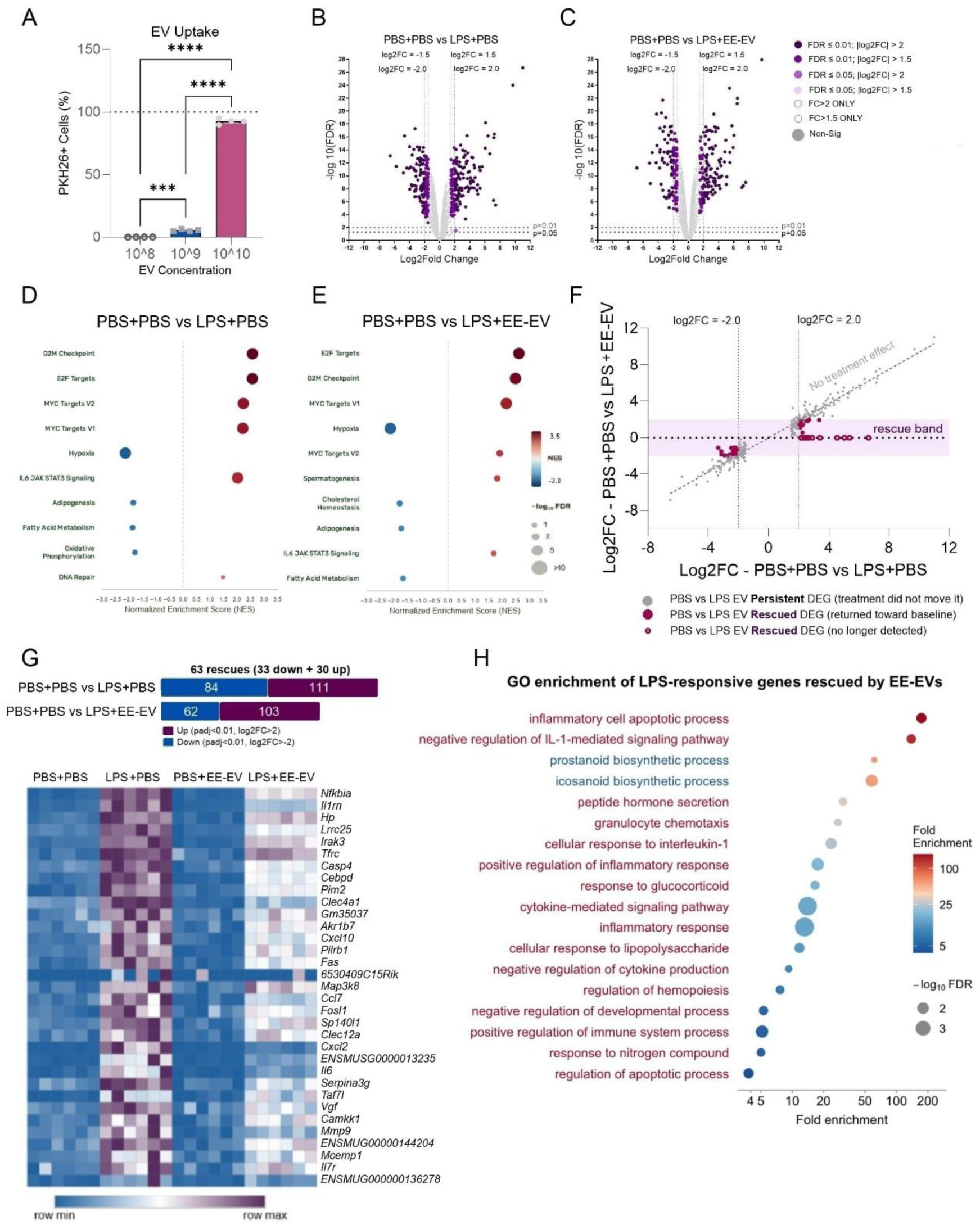
Milk-derived extracellular vesicles (EVs) from environmentally enriched (EE) donor dams attenuate LPS-induced transcriptional responses in BV2 microglia cells. **(A)** EV uptake by BV2 cells following exposure to 10^8^, 10^9^, or 10^10^ EV particles/mL (Data expressed as mean ± SEM, n = 4 wells/group. ***p <0.001, ****p <0.0001). Volcano plots showing the distribution of (**B**) 13,056 in PBS+PBS versus LPS+PBS-treated BV2 cells and (**C**) 13,022 genes in PBS+PBS versus LPS+EE-EV-treated BV2 cells. Gene expression is shown as log2 fold change (x-axis) versus −log10 (FDR) on the y-axis. Each dot represents a gene, with colors indicating different combinations of fold-change and FDR thresholds, as defined in the figure key. Functional pathway enrichment of genes identified in the PBS+PBS versus (**D**) LPS+PBS and (**E**) LPS+EE-EV comparisons, respectively. (**F**) Fold-change rescue plot comparing log2FC values for PBS+PBS versus LPS+PBS (x-axis) and PBS+PBS versus LPS+EE-EV (y-axis) for genes significantly altered by LPS. Movement toward y = 0 indicates attenuation of LPS-associated gene expression changes following EE-EV treatment. (**G**) Summary of genes demonstrating an expression pattern consistent with transcriptional rescue, including 33 DEGs upregulated in LPS+PBS and downregulated in LPS+EE-EV treatment and 30 DEGs suppressed in LPS+PBS and upregulated in LPS+EE-EV treatment; heatmap shows expression pattern of the rescued 33 gene set. (**H**) GO biological process enrichment among LPS-responsive genes normalized by EE-EV treatment. Overrepresented GO BP terms for the 33 genes upregulated by LPS and returned toward control levels by EE-EVs (red labels) and the 30 genes downregulated by LPS and similarly normalized (blue labels). Non-redundant terms are shown, defined as terms with no other enriched term among their GO descendants. Point position gives fold enrichment (log_10_ scale), point area gives the number of input genes annotated to the term, and fill gives −log_10_ FDR. All terms shown are FDR < 0.05 (Fisher’s exact test, Benjamini-Hochberg FDR correction). Fold enrichment for GO:0006925 and GO:2000660 exceeded the value reported by PANTHER (“>100”) and were calculated from reference counts. LPS: lipopolysaccharide; EE: environmental enrichment; EV: extracellular vesicles.

Following confirmation of EV uptake, RNA sequencing was used to characterize transcriptional responses of BV2 cells to LPS in the presence or absence of EVs (**Figure 3B-H**). LPS+PBS treatment resulted in 195 differentially expressed genes (DEGs) compared with PBS+PBS, including 111 upregulated and 84 downregulated genes (**Figure 3B**). As expected, LPS+PBS treatment resulted in a robust enrichment of genes involved in cytokine signaling, chemokine-mediated signaling, NF-ĸB signaling, innate and pattern recognition responses, and redox, metabolic, and cellular homeostasis (**Supplemental Figure 4**), consistent with activation of an inflammatory transcriptional response. Functional enrichment analysis further demonstrated significant enrichment of pathways associated with IL6/JAK/STAT3 signaling, E2F target genes, GM2 checkpoint, and MYC target gene programs (**Figure 3D**).

We next examined whether EE-EV treatment altered the transcriptional response of BV2 cells exposed to LPS. Comparison of PBS-treated cells with LPS+EE-EV treated cells identified 165 DEGs, consisting of 103 upregulated and 62 downregulated genes (**Figure 3C**). Pathway enrichment analysis of these DEGs revealed positive enrichment of IL6/JAK/STAT3 E2F targets, G2M checkpoint, MYC targets, and IL6/JAK/STAT3 signaling, with negative enrichment of hypoxia, cholesterol homeostasis, adipogenesis, and fatty acid metabolism (**Figure 3E**). Additional pairwise analyses were performed to distinguish transcriptional effects attributable to LPS, EE-EVs and their interaction (see **Supplemental Figure 5**). Direct comparison of LPS+PBS with LPS+EE-EV cells showed negative enrichment of IL6/JAK/STAT3, inflammatory response, TNFα signaling via NF-κB, and interferon responses following EE-EV treatment (**Supplemental Figure 5C-D**).These findings indicate that EE-EV treatment was associated with alterations in the transcriptional profile of LPS-exposed BV2 cells (see also **Supplemental Figure 4**).

To visualize whether EE-EVs attenuated LPS-associated changes in gene expression, we compared the log2FC between the PBS+PBS versus LPS+PBS and the PBS+PBS versus LPS+EE-EV comparisons (**Figure 3F**). Genes that moved toward a log2FC of zero following EE-EV treatment were considered to demonstrate an expression pattern consistent with transcriptional rescue. Overall, 63 genes demonstrated reversal of their LPS-associated expression changes, including 33 genes that were upregulated by LPS and subsequently reduced following EE-EV treatment and 30 genes that were downregulated by LPS and subsequently increased following EE-EV treatment (**Figure 3G**). Some representative inflammatory genes transcriptionally rescued by EE-EV treatment include *Il6*, *Il1rn*, *Cxcl10*, *Nfkbia*, *Irak3*, and *Slc7a11* (see **Supplemental Figure 4**), which are involved in cytokine and chemokine signaling, NF-κB regulation, innate immune function, and redox and homeostatic responses (Liu et al., 2012; Liu et al., 2017; Freihat et al., 2019; Ahsan et al., 2026; Nakazato et al., 2017; Lively et al., 2018; Sun et al., 1993; Wang et al., 2023; Jog et al., 2021; Bereshchenko et al., 2019; Ryan et al., 2017; Pinheiro et al., 2013; Lee et al., 2000; Vaillancourt et al., 2021; Peperzak et al., 2013; Lv et al., 2025; Vahsen et al., 2023; Könnecke et al., 2013; Yu et al., 2020; Uto et al., 2016; Hou et al., 2021; Qin et al., 2025; Wang et al., 2026). Heatmaps visually demonstrated the reversal of LPS-associated expression patterns across both rescued gene sets (**Figure 3G; Supplemental Figure 6**).

Gene ontology (GO) enrichment analysis of the rescued gene sets revealed distinct biological effects of EE-EV treatment (**Figure 3H**). The 33 DEGs induced by LPS and rescued (no longer DEG) by EE-EVs were strongly enriched for innate immune signaling. Cytokine-mediated signaling (9 genes, 14.0-fold, FDR 1.8 × 10⁻⁵) and inflammatory response (10 genes, 13.1-fold, FDR 1.0 × 10⁻⁵) were the most significant GO terms, with more specific enrichment for GO terms cellular responses to interleukin-1 (4 genes, 23.6-fold), lipopolysaccharide (5 genes, 11.7-fold), granulocyte chemotaxis, and both positive and negative regulation of inflammatory cytokine output. Response to glucocorticoid (4 genes, 16.6-fold) and peptide hormone secretion (3 genes, 30.7-fold) were also enriched.

In contrast, the 30 DEGs suppressed by LPS and increased by EE-EVs showed GO enrichment primarily for icosanoid and prostanoid biosynthesis (4 and 3 genes, 57.9- and 61.3-fold), reflecting a shared set of prostaglandin synthesis genes. Together, these findings indicate that EE-EVs act predominantly on the cytokine and pattern-recognition arm of the LPS response, with a narrower effect on icosanoid biosynthesis.

### Candidate milk-derived EE-EV miRNAs are associated with inflammatory and homeostatic gene expression patterns

Cross-referencing our previously-described milk-derived EE-EV miRNA cargos (Martz et al., 2026) against the present BV2 dataset identified candidate miRNA-mRNA relationships among the LPS-responsive genes that were normalized by EE-EV treatment (**Supplemental Table 2;** see also **Supplemental Figure 4**). Among these, mir-7b is a downregulated differentially expressed miRNA found in milk-derived EE-EVs that was associated with the predicted targets *Tsc22d3* (GILZ), and *Slc7a11*. These genes were no longer differentially expressed following EE-EV treatment in LPS exposed BV2 cells, consistent with normalization of an endogenous regulator of NF-ĸB signaling and redox homeostasis, respectively (Ahsan et al., 2026; Pinheiro et al., 2013; Liu et al., 2022; Bereshchenko et al., 2019; André et al., 2016; Wang et al., 2026). Following a similar pattern, miR-132-3p was associated with normalization of *Slc13a3*, suggesting another potential association with redox/cellular homeostasis (Zhao et al., 2024; Strungaru et al., 2011). In contrast, upregulation of the differentially expressed miR-532-3p was associated with EE-EV-driven reversal of LPS-induced *Il1rn* in BV2 cells, mechanistically linking milk-derived EE-EV miRNA cargos to inflammatory modulation (Lively et al., 2018).

## Discussion

Early parental experiences can influence offspring through multiple interacting pathways that converge on various physiological and molecular signals transmitted across development (Flemming et al., 1999; Curley and Champagne, 2016; Rincón-Cortés, 2024; Kosie et al., 2024; de Weerth et al., 2023). Among these pathways, our findings support milk-derived EVs as a putative biological route through which parental experiences may be communicated to the developing offspring via nursing (see summary **Figure 4**). Administration of EE programmed milk-derived EVs was sufficient to ameliorate deficits in social behavior in MIA-exposed rat offspring. Moreover, EE-EVs promoted social behavior in rat pups from unchallenged litters, consistent with the literature that EE exposure augments sociability (Núñez Estevez et al., 2020; DeRosa et al., 2022; Kentner et al., 2016; Martz et al., 2026; Zhao et al., 2021; D’Andrea et al., 2007). EE-EV treatment also attenuated LPS-induced transcriptional responses in BV2 microglia. Together with our previous findings that maternal EE remodels milk-derived EV miRNA cargo and attenuates MIA-associated molecular changes in offspring brain and behavior (Martz et al., 2026), and prior results linking microglia to social deficits in MIA model offspring (Ikezu et al., 2021; Kalavai et al., 2021), the present data suggest that environmental programming of milk-derived EVs may have functional consequences for offspring development. Encouragingly, these findings also support the idea that the parental environment does not simply transmit developmental risk but can also provide molecular signals that may promote resilience.

**Figure 4.**
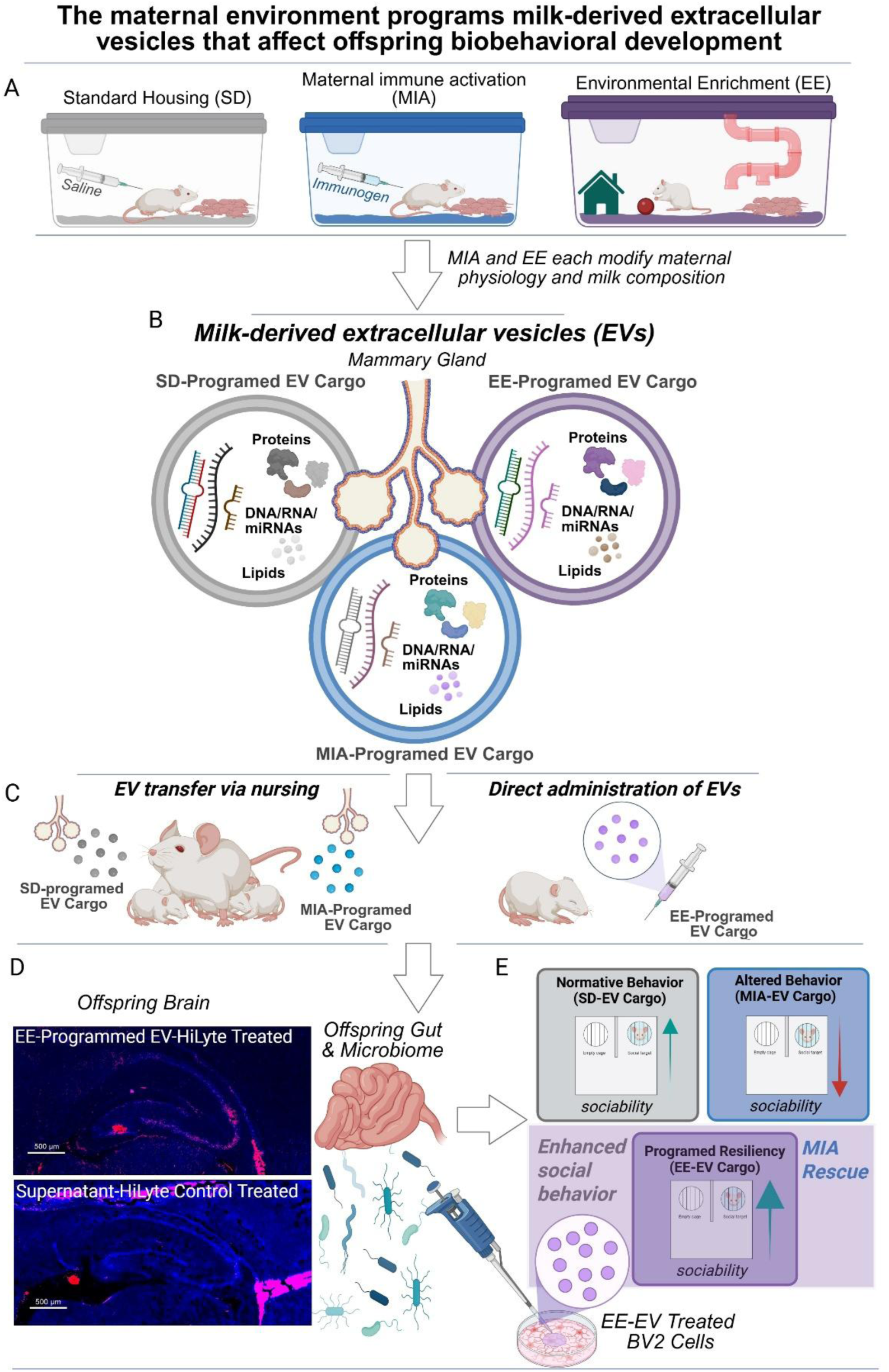
Proposed model linking maternal environmental experience, milk-derived extracellular vesicles, and offspring neuroimmune and behavioral outcomes. **(A)** Maternal experimental conditions: standard laboratory housing (SD), maternal immune activation (MIA), and environmental enrichment (EE) housing. **(B)** Maternal environmental experience serves as a cue to program the molecular cargo of milk-derived extracellular vesicles (EVs), including miRNAs, resulting in distinct EV populations associated with each condition. **(C)** Milk-derived EVs and their cargo are transferred naturally to offspring through nursing or experimentally administered directly to offspring. Following exposure, **(D)** EVs may interact with peripheral tissues and the brain, sending programming signals (e.g., miRNAs) that can shape development. Representative images show hippocampal localization of fluorescently labeled EE-programmed EVs, whereas depleted supernatant controls show no detectable hippocampal fluorescence. Potential interactions among the gut, microbiome, other peripheral tissues, and brain are depicted as pathways linking EV exposure to offspring neurodevelopment and behavior. **(E)** MIA was associated with reduced sociability, whereas administration of EE-programmed EVs to MIA-exposed offspring restored social behavior. Neonatal administration of EE-EVs also promoted social behavior in healthy unchallenged rats. In BV2 microglial cells, EE-EVs attenuated LPS-associated inflammatory transcriptional responses and modulated metabolic and cellular homeostatic pathways, supporting a potential neuroimmune mechanism underlying these behavioral effects. LPS: lipopolysaccharide; EE: environmental enrichment; EV: extracellular vesicles. *Created in BioRender. Kentner, A. (2026)* https://BioRender.com/p1vpez6.

The potential relevance of milk as a route of biological communication is supported by evidence showing that it contains bioactive hormones, such as gonadotropin releasing hormone (Baram et al., 1977), illustrating that lactation can convey signaling molecules in addition to nutritional substrates. More recently, milk-derived EVs have emerged as particularly interesting candidates because they package diverse regulatory molecules within membrane-enclosed vesicles, putatively protecting and delivering this cargo to recipient cells (Welsh et al., 2024; Jiang et al., 2021; Kalluri et al., 2020). Milk-derived EVs have also been detected in the developing brain following maternal nursing and experimental oral administration, where they have crossed brain endothelial barriers and accumulated in regions like the hippocampus (Manca et al., 2018; López de Las Hazas et al., 2022; Zhou et al., 2022; Haney et al., 2019). In this context, the present behavioral findings add to a growing body of evidence (Li et al., 2026; Zhou et al., 2022) suggesting that milk-derived EVs can influence offspring phenotype, rather than merely serve as biomarkers of maternal experience.

The behavioral effects of EE-EVs were most evident during juvenile development in our studies. MIA-associated reductions in social investigation were prevented in male offspring and were not secondary to generalized alterations in activity or anxiety-like behavior. Female offspring displayed a related effect on social play vocalizations, highlighting the sex- and behavioral-specificity components of the response. The observation that EE-EVs also increased social investigation in healthy juvenile males is particularly notable because it suggests that these EVs may not simply normalize a MIA-induced phenotype but may actively promote aspects of social development. This is consistent with previous work demonstrating that the effects of maternal EE on offspring sociability can depend on the lactational environment. Specifically, evidence using cross-fostering methods identified a postnatal maternal contribution to enhanced social behavior in male offspring (DeRosa et al., 2022). Combined, these findings support the possibility that the influence of the lactational period is mediated by molecular components of milk, including EV cargo (Jiang et al., 2021; Kalluri et al., 2020). However, whether early-versus later lactational EV populations differ in their biological effects remains an important question. In this present proof-of-concept study, EVs were pooled across the milk collection period to reflect the integrated population of vesicles to which nursing offspring are exposed; future studies will need to determine whether temporally diverse EV populations exert discrete or overlapping developmental effects.

Given the relationship between MIA and the mediating effects of microglia on sociability (Ikezu et al., 2021; Kalavai et al., 2021), a possible cellular mechanism emerged from our studies of BV2 microglia. Milk-derived EVs were internalized by BV2 cells and EE-EVs shifted a subset of LPS-responsive genes toward the unstimulated state, including *lL6*, *Cxcl10*, *Nfkbia*, *Irak3*, *Il1rn*, and *Casp4*. These genes span both inflammatory effectors and endogenous feedback mechanisms that are engaged during innate immune activation. For example, *Il6* promotes inflammatory signaling, whereas *Nfkbia* encodes IĸBα, a canonical negative regulator of NF-ĸB signaling. Moreover, IRAK3/IRAK-M limits TLR/IL-1 receptor signaling upstream of NF-ĸB activation and IL-1 receptor antagonist (IL-1Ra; *il1rn*) provides an additional endogenous brake on inflammatory signaling (Chatterjee et al., 2017; Kobayashi et al., 2002; Freihat et al., 2019; Lamacchia et al., 2010; Sun et al., 1993; Yu et al., 2020).

LPS- and stress-induced reductions in social behavior have been linked previously to IL-1 signaling and can be attenuated by IL-1 receptor antagonist (IL1Ra; Arakawa et al., 2009; Konsman et al., 2008). In BV2 cells, LPS increased *Il1rn* expression, which encodes IL-1Ra, whereas EE-EVs reduced this response. Combined with our other gene expression data, normalization of *Il1rn* may reflect reduced upstream inflammatory drive and need for compensatory IL-1 signaling. Importantly, these data reflect *Il1rn* transcription and not changes in IL1Ra protein or signaling activity. Together, these findings identify altered neuroinflammatory regulation as a potential pathway linking EE-EV exposure to behavioral rescue by EE-EVs. This is in line with other studies demonstrating that milk-derived EVs are taken up by microglia, which can alter their cellular responses (Akinduro et al., 2025; Storm et al., 2025; Wijenayake et al., 2024; Li et al., 2026).

Integration of the BV2 transcriptome with our previously identified EE-EV miRNA cargo (Martz et al., 2026) further identified candidate molecular links. Notably, increased milk-derived EV miR-532-3p was associated with normalization of *Il1rn*, while reduced miR-7b was associated with normalization of *Tsc22d3*. *Tsc22d3 encodes* GILZ, a glucocorticoid-responsiveness regulator with established anti-inflammatory effects through the modulation of NF-ĸB and related pathways (Bereshchenko et al., 2019; André et al., 2016; Pinheiro et al., 2013).

Although not a significantly differentially expressed miRNA, miR-23a-3p was present in our milk-derived EE-EVs (Martz et al., 2026) and was associated with normalization of *Fas* and *Chst7*, which were LPS-responsive and rescued by EE-EVs in BV2 cells. In a recent study, blood-derived EV-associated miR-23a-3p increased sociability in mice predisposed to social deficits (Matoba et al., 2026), providing additional evidence that EV miRNAs have functional effects. This treatment did not increase sociability in wild-type mice (Matoba et al., 2026), in line with the idea that vulnerabilities/early life challenges may make organisms more receptive or sensitive to context and to supportive resources (Boyce and Ellis, 2005). Our findings extend these prior findings by showing that milk-derived EVs programmed by maternal EE promotes offspring social behavior in both natural and pathological contexts and links EV-mediated behavioral effects to the transmission of the maternal environment. Moreover, these observations highlight that the effects of EVs on behavior may depend on a combination of the EV source, cargo, and recipient.

The miRNA-mRNA relationships identified here are predicted, not validated, and miRNA is only one class of many bioactive molecule cargos in EVs (Welsh et al., 2024; Jiang et al., 2021). Despite these limits, however, the convergence EV miRNA cargo, transcriptional normalization, and behavior we report here supports a functionally relevant role for milk-derived EVs in development and MIA rescue. This interpretation is further supported by our previous demonstration that EE alters the miRNA cargo of milk-derived EVs compared to both standard laboratory housed and MIA-exposed groups (Martz et al., 2026). Together, these findings position milk-derived EVs as a potential molecular pathway through which maternal environmental experience is transmitted to the developing offspring.

## Conclusions

Overall, this work shows that supportive parental environments impact social development not only through caregiving behaviors, but also through EV-mediated molecular signals transmitted through nursing. Defining the specific EV cargo that mediates these effects will be central to understanding how maternal environmental experience is encoded in milk-derived EVs to influence offspring.

It is important also to keep in mind the broader biological and social context in which lactation occurs, including the conditions that support breastfeeding people and their families. At the same time, lactation is only one of many pathways through which parental experiences may be communicated to offspring. The goal is not to define a single route to healthy development, but to understand how milk, caregiving, parental physiology, and other components of the family environment can interact to shape the molecular signals and developmental conditions that promote resilience.

## Supporting information

Supplemental Table 1. MIA Reporting Table

## Author Contributions

GDASB, PP, TTQL, and ACK performed milk collection, extracellular vesicle (EV) validation/characterization, and/or EV administration. GDASB, PP, HV, TTQL, and ACK ran behavioral experiments and analyzed the resulting data; ACK made the associated figures. GDASB and PP contributed the fluorescently tagged EV and control images shown in Figure 4. HH and YA carried out the BV2 microglial experiments, SB, GDASB, and ACK analyzed the BV2 sequencing data; ACK made the associated figure. ACK wrote the manuscript and GDASB, PP, HV, HH, YA, TTQL, SB, and ACK contributed to manuscript editing. SB and ACK were involved in funding acquisition, project design, and supervised trainees. ACK conceptualized and directed the project. All authors approved the final version of the manuscript.

## Data Availability

All data is presented in the article files or available upon request.

## Code Availability

The code is available on request.

---

**Supplemental Figure 1.**
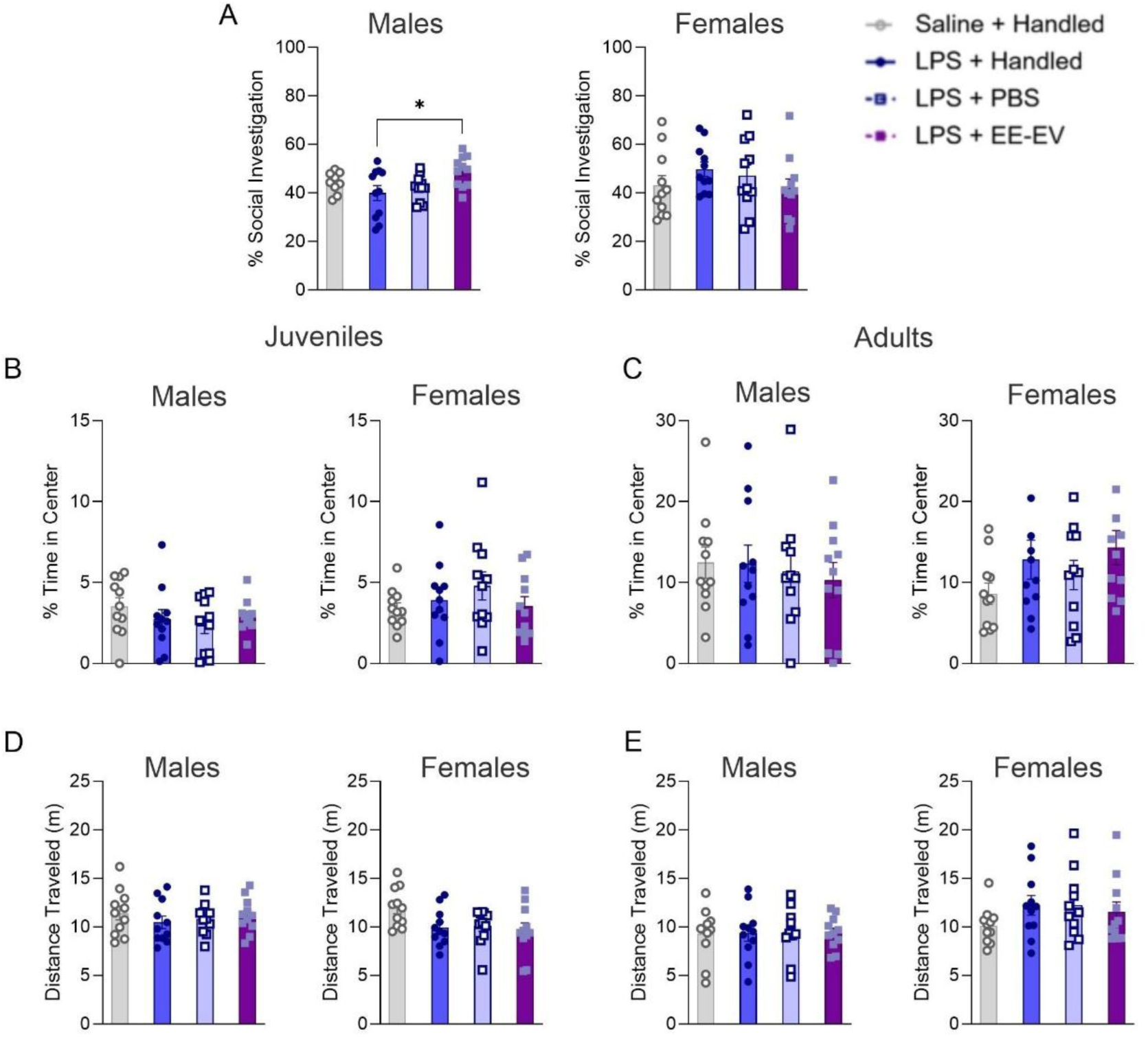
Milk-derived extracellular vesicles (EVs) from environmentally enriched (EE) donor dams increase adult offspring social behavior. Adult male and female offspring behavioral outcomes for **(A)** percent time investigating a novel conspecific. Percent time spent in the center of the open field and total distance traveled (m) for male and female juvenile **(B, D)** and adult **(C, E)** offspring. Data expressed as mean ± SEM, n = 11-12 litters/group. *p <0.05, **p <0.01, ***p <0.001, ****p <0.0001. LPS: lipopolysaccharide; EE: environmental enrichment; EV: extracellular vesicles.

**Supplemental Figure 2.**
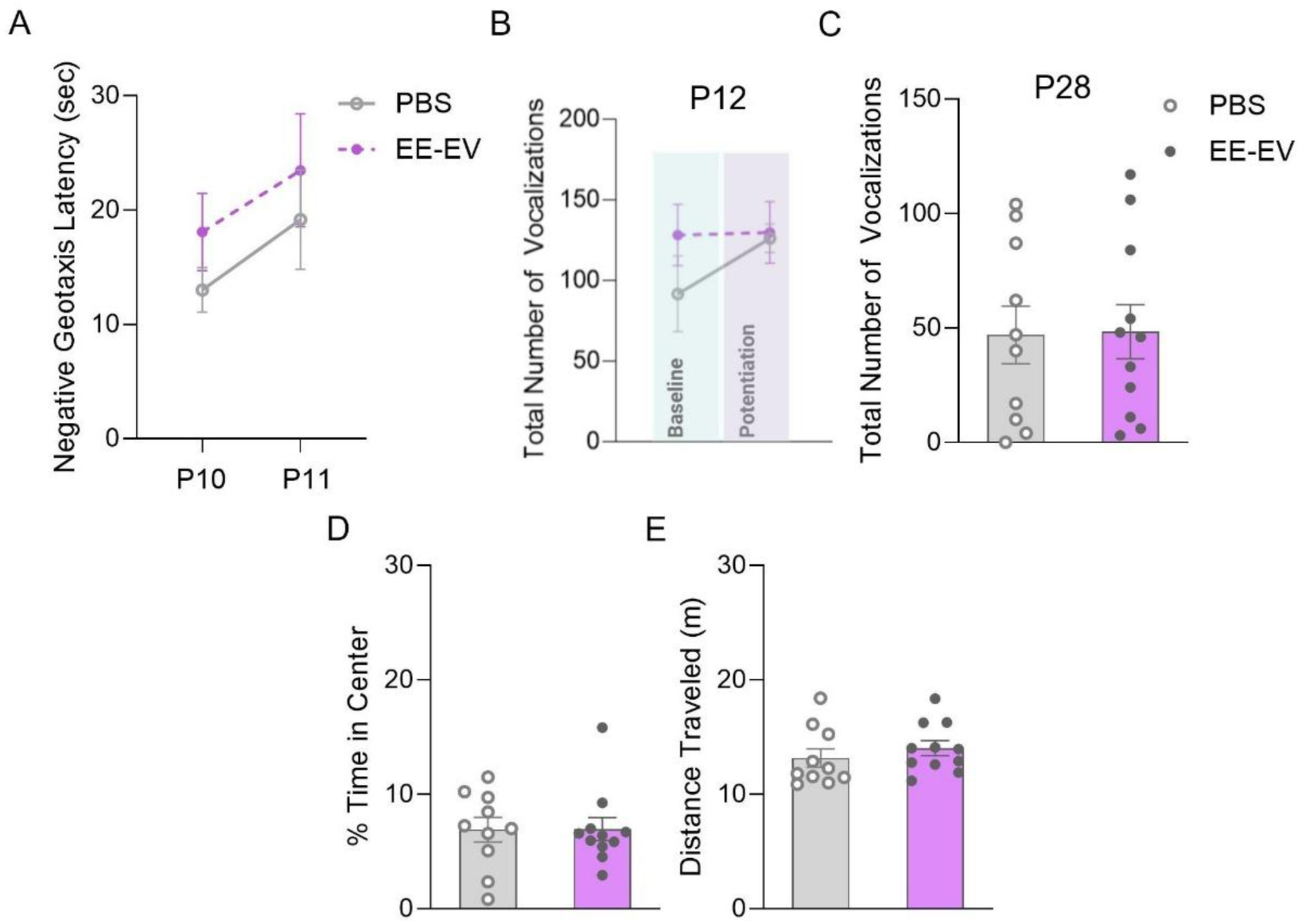
Milk-derived extracellular vesicles (EVs) from environmentally enriched (EE) donor dams do not affect anxiety-like or locomotor behavior in healthy unchallenged offspring. Male behavioral outcomes for **(A)** neonatal negative geotaxis latency, total number of **(B)** P12 isolation-induced and **(C)** P28 social play vocalizations emitted, and open field behavior for **(D)** percent time spent in the center and **(E)** distance traveled (meters) in the arena. Data expressed as mean ± SEM, n = 10-11 litters/group. *p <0.05, **p <0.01, ***p <0.001, ****p <0.0001. EE: environmental enrichment; EV: extracellular vesicles.

**Supplemental Figure 3.**
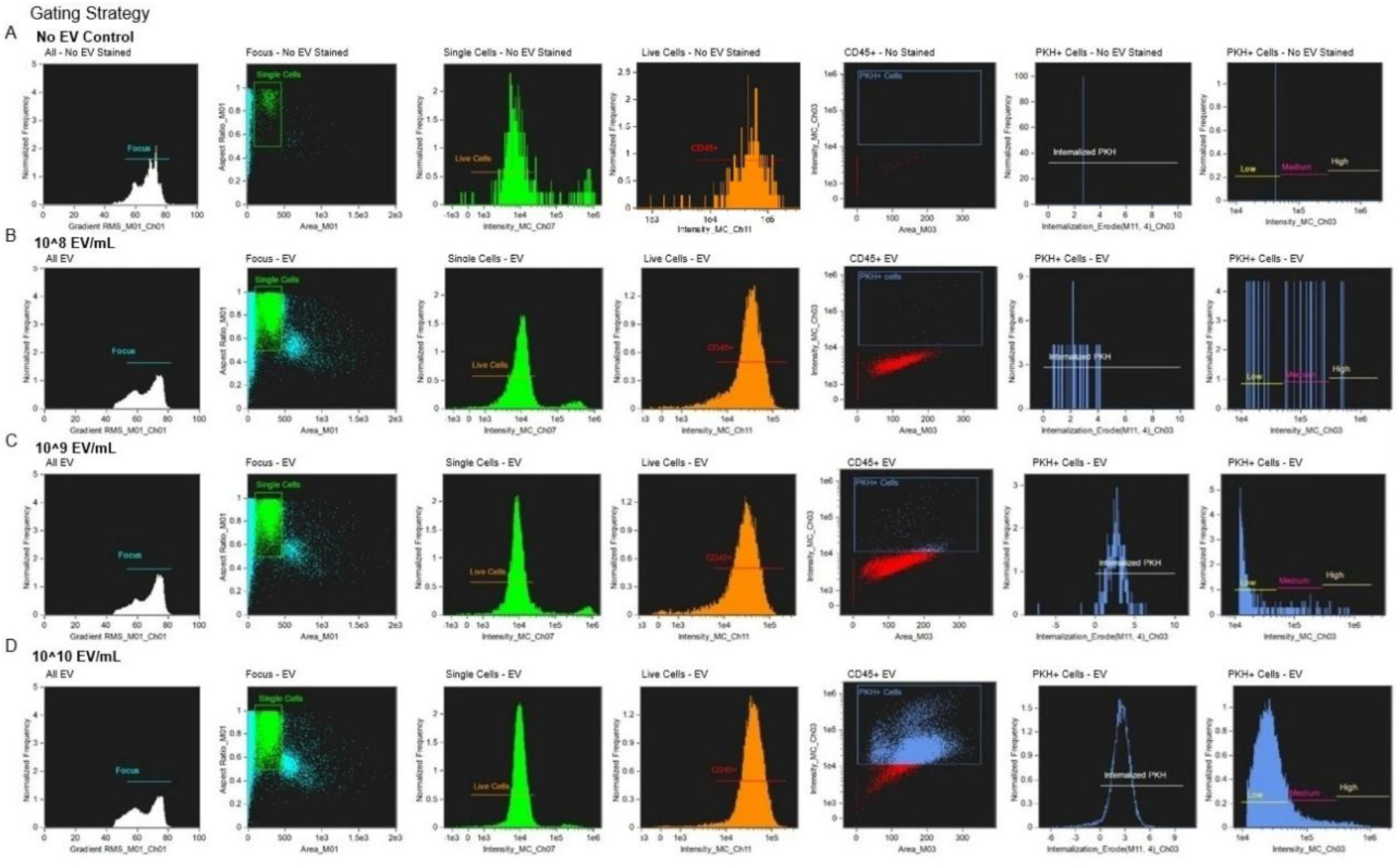
ImageStream gating strategy for assessment of milk-derived extracellular vesicle (EV) uptake by BV2 cells. Representative ImageStream plots demonstrating the sequential gating strategy used to identify PKH-26 (PKH)-labeled EV-positive BV2 cells. Representative plots are shown for (**A**) BV2 cells without PKH EV, and BV2 cells exposed to (**B**) 10^8^, (**C**) 10^9^, (**D**) 10^10^, particles/mL of PKH EV. For each condition, the sequential gates identify all events, focused cells, single cells, live cells, CD45+ cells, and PKH+ cells, with the final PKH+ population assessed for EV internalization and fluorescence intensity. The sample without EV was used to establish gating thresholds for identification of PKH+ cells.

**Supplemental Figure 4.**
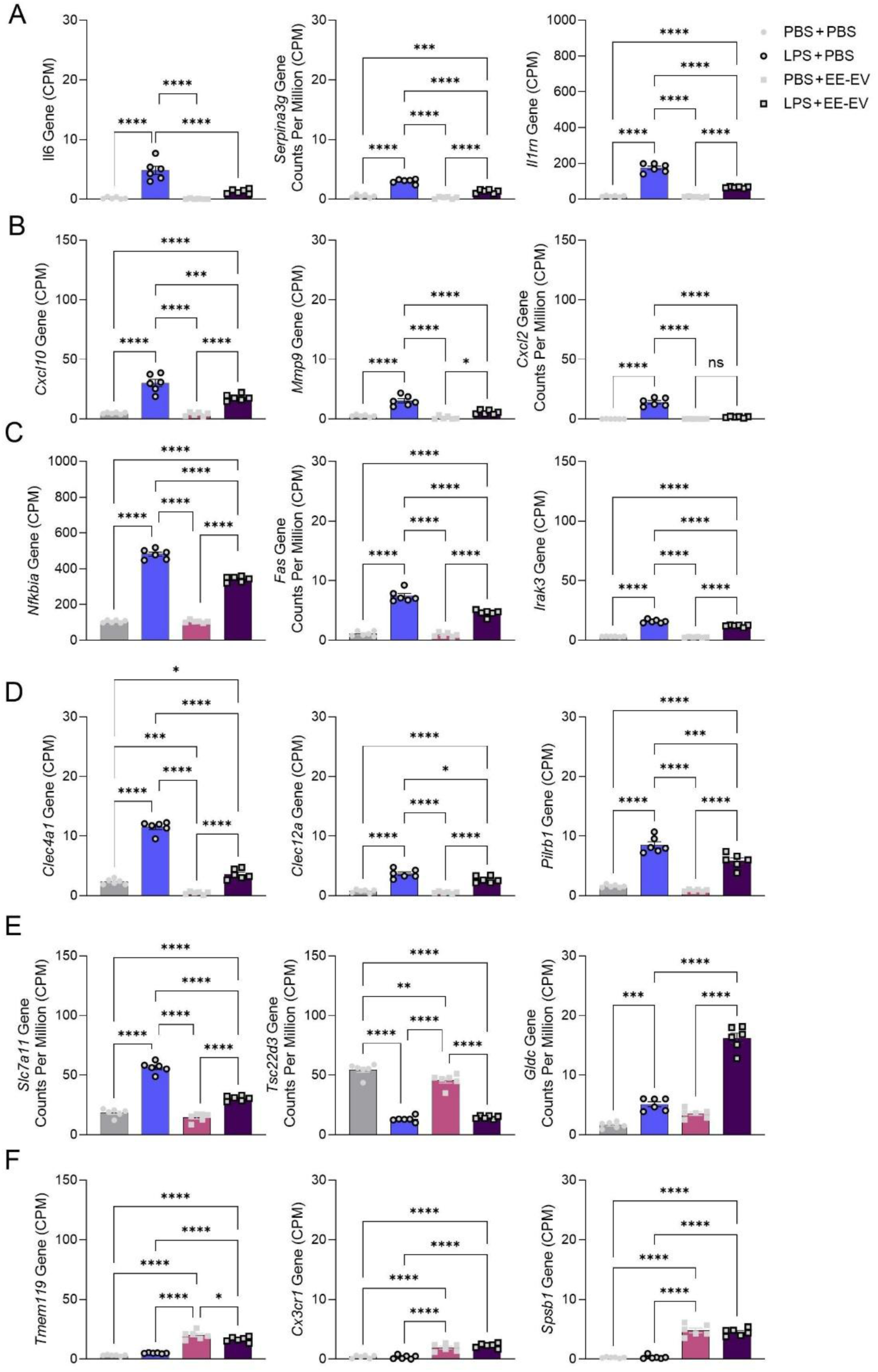
Milk-derived extracellular vesicles (EVs) from environmentally enriched (EE) donor dams reverses representative LPS-responsive inflammatory gene expression in BV2 cells. Gene expression is shown as counts per million (CPM) for representative genes associated with broad categories of **(A)** cytokine signaling and inflammatory mediators, **(B)** chemokine and extracellular inflammatory signaling, **(C)** NF-ĸB signaling, **(D)** innate immune receptors and immunoregulatory signaling, and **(E)** redox, metabolic, and cellular homeostasis. In addition to reversing LPS-associated inflammatory gene expression, EE-EVs induced a distinct transcriptional response in BV2 cells. **(F)** Representative genes (Tmem119, Cxcr1, and Spsb1) demonstrating an EE-EV-associated transcriptional response independent of LPS treatment; these genes were not significantly altered by LPS but were significantly increased following EE-EV treatment. Data expressed as mean ± SEM, n = 6 wells/group*p <0.05, **p <0.01; ***p <0.001. LPS: lipopolysaccharide; EE: environmental enrichment; EV: extracellular vesicles.

**Supplemental Figure 5.**
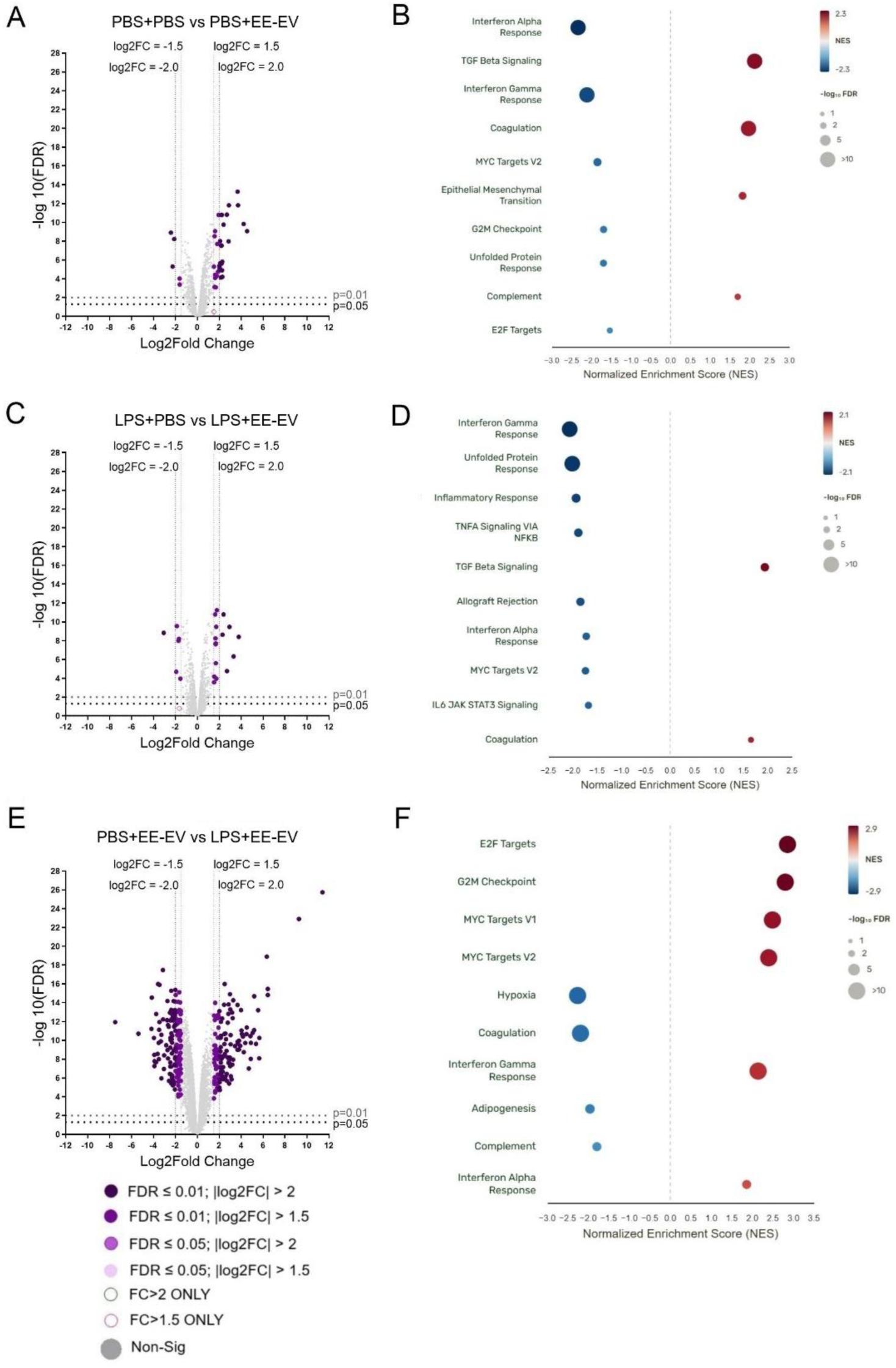
Differential gene expression and functional enrichment across additional BV2 treatment comparisons. Volcano plots and functional pathway enrichment of genes identified in the pairwise comparisons of (**A, B**) PBS+PBS versus PBS+EE-EV (**C, D**) LPS+PBS versus LPS+EE-EV, and (**E, F**) PBS+EE-EV versus LPS+EE-EV. Gene expression is shown as log2 fold change (x-axis) versus −log10 (FDR) on the y-axis. Each dot represents a gene, with colors indicating different combinations of fold-change and FDR thresholds, as defined in the figure key. Functional enrichment plots show the biological pathways and processes associated with differentially expressed genes identified in each comparison. BV2 cells were treated with PBS or LPS in the presence or absence of EE-EVs. LPS: lipopolysaccharide; EE: environmental enrichment; EV: extracellular vesicles.

**Supplementary Figure 6.**
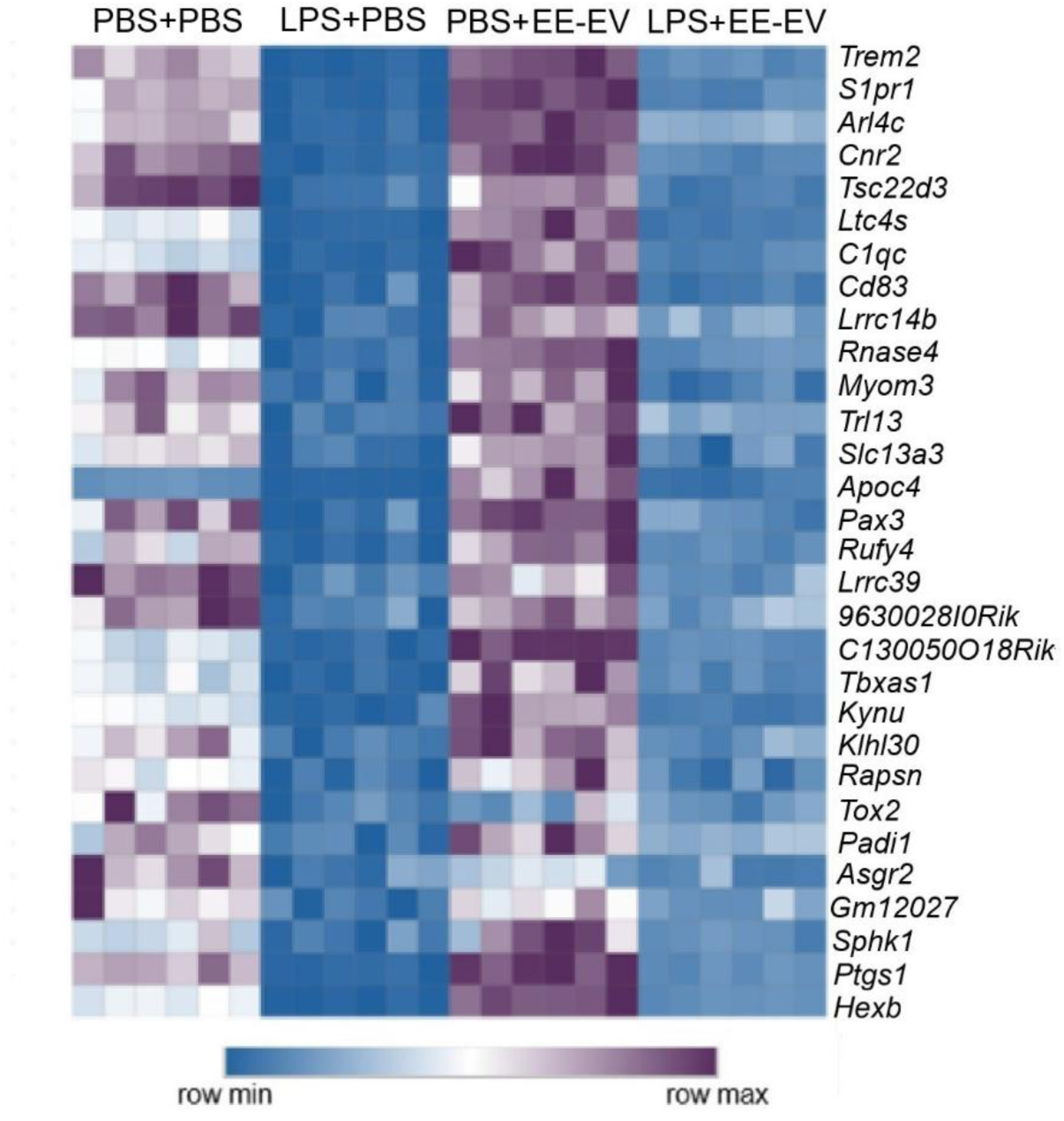
Transcriptional rescue of LPS-downregulated genes in BV2 microglial cells by milk-derived extracellular vesicles (EVs) from environmentally enriched (EE) donor dams. Heatmap showing the 30 differentially expressed genes (DEGs) downregulated in LPS+PBS BV2 cells and subsequently upregulated following LPS+EE-EV treatment. Rows represent genes and columns represent individual samples, illustrating the reversal of LPS-associated gene expression following EE-EV treatment. Genes were identified as differentially expressed (DEG) when they demonstrated an absolute log2 fold change (|log2FC|) ≥2 and an FDR ≤0.01. LPS: lipopolysaccharide; EE: environmental enrichment; EV: extracellular vesicles.

**Supplemental Table 1. Maternal Immune Activation (MIA) Model Reporting Guidelines Checklist.** See accompanying PDF file.

**Supplemental Table 2.**
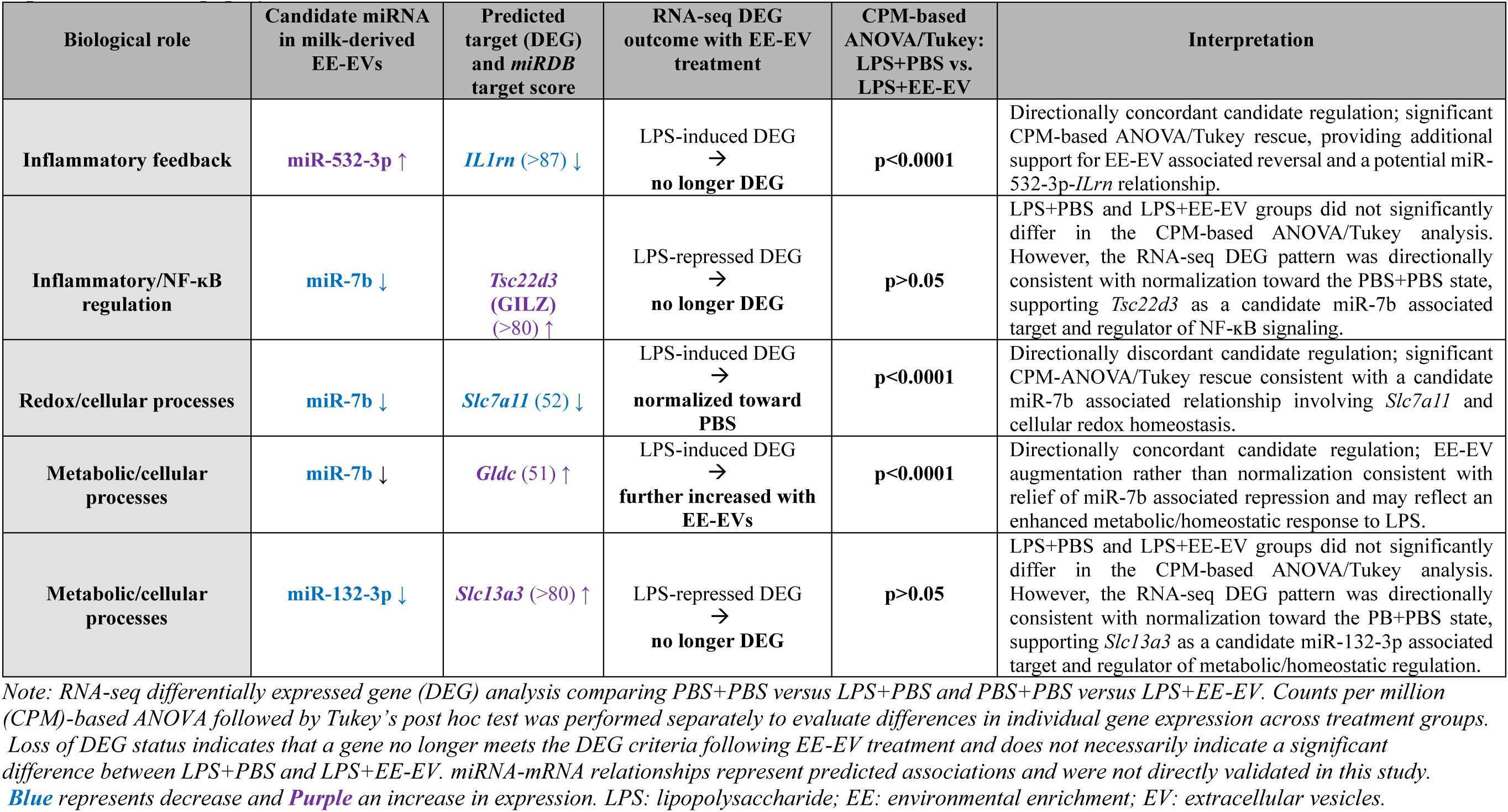
Candidate milk-derived EE-EV miRNA–mRNA relationships associated with normalization of LPS-responsive gene expression. LPS: lipopolysaccharide; EE: environmental enrichment; EV: extracellular vesicles.

